# Scrutinizing surface glycoproteins and poxin-schlafen protein to design a heterologous recombinant vaccine against monkeypox virus

**DOI:** 10.1101/2020.01.25.919332

**Authors:** Syeda Farjana Hoque, Md. Nazmul Islam Bappy, Anjum Taiebah Chowdhury, Md. Sorwer Alam Parvez, Foeaz Ahmed, Md. Abdus Shukur Imran, Kazi Faizul Azim, Mahmudul Hasan

**Affiliations:** Faculty of Biotechnology and Genetic Engineering, Sylhet Agricultural University, Sylhet-3100, Bangladesh; Department of Pharmaceuticals and Industrial Biotechnology, Sylhet Agricultural University, Sylhet-3100, Bangladesh; Department of Genetic Engineering and Biotechnology, Shahjalal University of Science and Technology, Sylhet-3114, Bangladesh; Department of Microbial Biotechnology, Sylhet Agricultural University, Sylhet-3100, Bangladesh

**Keywords:** Monkeypox, recombinant vaccine, surface binding protein, poxin-schlafen protein, envelope protein

## Abstract

Monkeypox is a zoonotic disease caused by monkeypox virus with noteworthy mortality and morbidity. Several recent outbreaks and the need of dependable reconnaissance have raised the level of concern for this developing zoonosis. In the present study, a reverse vaccinology strategy was developed to construct a peptide vaccine against monkeypox virus by exploring cell surface binding protein, Poxin-Schlafen andenvelope protein. Both humoral and cell mediated immunity induction were the main concerned properties for the designed peptide vaccine. Therefore, both T cell and B cell immunity against monkeypox virus were analyzed from the conserver region of the selected protein. Antigenicity testing, transmembrane topology screening, allergenicity and toxicity assessment, population coverage analysis and molecular docking approach were used to create the superior epitopes of moneypox virus. The subunit vaccine was constructed using highly immunogenic epitopes with appropriate adjuvant and linkers. Molecular docking examination of the refined vaccine with various MHCs and human immune receptor illustrated higher binding interaction. The designed construct was reverse transcribed and adjusted for *E. coli* strain K12 earlier to inclusion inside pET28a(+) vector for its heterologous cloning and expression. The study could start in vitro and in vivo studies concerning effective vaccine development against monkeypox virus.

## Introduction

The destruction of smallpox was one of the most noteworthy accomplishments within the history of public wellbeing. One of the awful incongruities of this victory is the rise of Monkeypox (MPX) with noteworthy mortality and morbidity (Ladnyj et al., 1972). The later clear increment in human monkeypox cases over a wide geographic range, the potential for advance spread, and the need of dependable reconnaissance have raised the level of concern for this developing zoonosis (Durski et al., 2018). Monkeypox could be a smallpox-like viral contamination caused by an infection of zoonotic beginning, which has a place to the sort Orthopoxvirus, family Poxviridae, and sub-family Chordopoxvirinae.

Monkeypox infection could be a large and double stranded DNA virus (Damon, 2011). The symptoms of the monkeypox are similar like the clinical symtoms of smallpox. It contain various types of clinical presentation which include fever, back pain,characteristic rash, headache and flulike symptoms (Pal et al., 2017).Transmission of monkeypox is accepted to happen by means of respiratory excretions or outside fabric or contact with injury exudate. Viral shedding through feces may speak to another introduction source (Jezek et al., 1988; Hutson et al., 2009). Human monkeypox was first found in Congo in 1970 (Arita et al., 1985; Marennikova et al., 1972). Human monkeypox may be a sporadic illness which was caused by MPV transmission from creatures to people within the tropical rainforest districts of Central and Western Africa and it was examined from 1981 to 1986. Approximately 28% of the cases were caused by secondary human to human spread. On the other hand, tertiary and quaternary transmissions of virus were uncommon (Ježek et al., 1988). Almost 388 of the 418 recorded cases of human monkeypox happened in Zaire from 1970 to 1995 (Breman, 2000). Human monkeypox cases were surpassed and it was 500 in 1996 and 1997. A few hundred modern cases were found by DRC wellbeing specialistsin January 1999 (Mukinda et al., 1997; Breman, 2000).

The suspension of general vaccination within the 1980s has given rise to expanding defenselessness to monkeypox infection within the human. Monkeypox has continuously been considered an uncommon intermittent malady with a restricted capacity to spread between people (WHO, 1984). It was one of the life-threatening illness within the Law based Republic of the Congo (DRC) and other nations of western and central Africa (Meyer et al., 2002). The recognizable proof of monkeypox in 3 isolated patients within the United Kingdom (UK) in September raised media and political consideration on a rising open wellbeing risk. As of 13th October 2018, there have been 116 affirmed cases, the larger part of which was beneath forty years old (Petersen et al., 2019). The risk would increment in the event that there would be a destructiveness increment (Blumberg and Lloyd-Smith, 2013; Jackson et al., 2001), a infection spill into more broadly conveyed taxa (Reynolds et al., 2012) or an presentation in other landmasses (Rimoin et al., 2010).Thus, monkeypox has a place to the biosafety level 3, the “high threat” biodefence level within the EU (Tian et al., 2014).The mortality rate of monkeypox (10%) lies between the mortality rate of variola minor(1%) and variola major (30%). Monkeypox is an endemic disease in the Democratic Republic of the Congo. The virus was also imported once into the USA (Sklenovská et al., 2018).

There are no authorized treatments to treat human monkeypox virus untill 2019.Dryvax, a smallpox vaccine were used for both smallpox and monkeypox treatment (Hammarlund et al., 2005). Both vaccine and persons in contact with the vaccine were influenced by the various negative effects (CDC, 2001; CDC, 2003). Monkeypox infection disease of sound rhesus macaques shows up to be a reasonable demonstrate to think about defensive resistant reactions against monkeypox (Hooper et al., 2004). Undoubtedly, macaques inoculated with Dryvax are ensured from monkeypox (Edghill-Smith et al., 2003; Earl et al., 2004; Hooper et al., 2004; Edghill-Smith et al., 2005).Later information have appeared that the major mode of security from monkeypox managed by the current non-attenuated smallpox immunization is intervened by Abs (Edghill-Smith et al., 2005). Exhaustion of either CD4 T cells or CD8 T cells in inoculated creatures some time recently monkeypox infection challenge does not influence survival, though B cell exhaustion some time recently and amid immunization abrogates vaccine-induced assurance. In like manner, passive administration of vaccinia infection (VACV) 3 Abs confers assurance from subsequent deadly monkeypox (Edghill-Smith et al., 2005). Hence, next-generation smallpox vaccination may not ought to be based on duplicating vectors, given that they evoke appropriate Ab reactions. The definition of VACV defensive Ags has been constrained by the complexity of the VACV proteome that encodes a few 200 proteins. In any case, proteins L1R and A27L, particular to the intracellular develop infection (IMV), and proteins A33R and B5R, particular to the extracellular encompassed infection (EEV), have been appeared to be immunogenic and ensured mice from VACV (Fogg et al., 2004; Galmiche et al., 1999; Hooper et al., 2000). As of late, security from monkeypox-induced extreme infection was too watched taking after gene gun immunizations with as it were four VACV DNA plasmids encoding these four proteins (Hooper et al., 2004). In this way, smallpox vaccines based on recombinant peptides ought to be able to bestow security from monkeypox.

Hence, it is necessary to invent the suitable polyvalent vaccine against monkeypoxvirus. Genomics and proteomics study of drug has a great wing to design drugs by utilizing in-silico study. In-silico study is also necessary for immunoinformatics approach, reducing cost and time in development process of drug (Hasan et al., 2019a; Adhikari et al., 2018; Mohammed et al., 2017; Dash et al., 2017). Different genome based innovations have empowered practically dazzle choice of vaccine candidates and permitted expectation of antigens without the necessity to develop the pathogen in vitro (Rappuoli, 2000; Sette and Fikes, 2003; Agrawal and Raghava, 2018). Subsequently, we in this connected a few bioinformatics database, tools and computer program to plan a thermostable and profoundly immunogenic, multi-epitope peptide vaccine against Monkeypox virus utilizing their preserved locale protein arrangements.

## Materials and Methods

In the present study, reverse vaccinology approach was used to design a novel multiepitope subunit vaccine against monkeypox virus requiring effective medications and preventive measures. The flow chart summarizing the entire protocol of in silico strategy for developing a vaccine has been illustrated in Figure 01.

**Figure 01:**
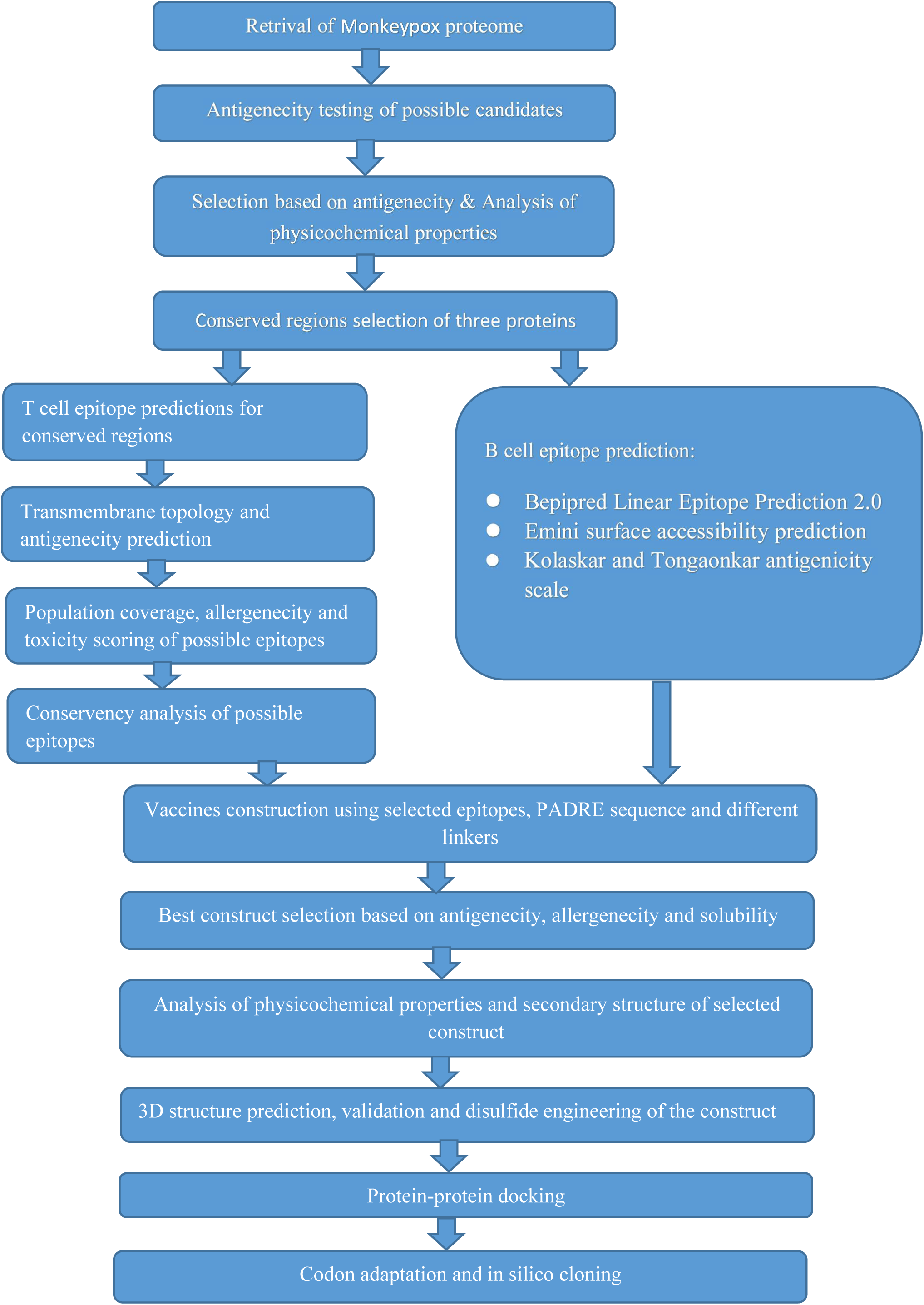
Schematic representation of the process used for epitope based vaccine developent against monkeypox virus.

### 1. Retrieval of protein sequences and antigenicity screening

The ViralZone (https://viralzone.expasy.org/search?query=monkeypox) and National Center for Biotechnology Information (NCBI) server were used for the comprehensive study and selection of monkeypox virus proteins (https://www.ncbi.nlm.nih.gov/genome/?term=monkeypox). Three proteins (Cell surface binding protein, Poxin-Schlafen, Envelope protein A28 homolog) were selected on the basis of their antigenicity by using VaxiJen v2.0 (http://www.ddg-pharmfac.net/vaxijen/) (Doytchinova and Flower, 2007a). ProtParam tool was used for the analysis of their various types of physicochemical properties (Gasteiger et al., 2003).

### 2. Identification of homologous proteins sets

Homologous sequences of the antigenic proteins (i.e. Cell surface-binding protein, Poxin-Schlafen, Envelope protein A28 homolog) in different monkeypox strain were retrieved from the NCBI database by using BLASTp tool. Proteins were used as query and the searches were restricted to Monkeypox virus (taxid:10244), Monkeypox virus Zaire-96-I-16 (taxid:619591), Monkeypox virus (strain Sierra Leone 70-0266) (taxid:130669), Monkeypox virus (strain Zaire 77-0666) (taxid:130670).

### 3. Identification of conserved regions and T-cell epitope prediction

Clustal Omega server was used for Multiple sequence alignment (MSA) (Chenna et al., 2003; Sievers and Higgins, 2014), as well as encourage those common locales were utilized for T-Cell epitopes expectation (Vita et al., 2014). T cell epitope prediction server (http://tools.iedb.org/main/tcell/) of IEDB were utilized to predict the T cell epitope (MHC-I and MHC-II).

### 4. Transmembrane topology and antigenicity screening of T-Cell epitope

Prediction of transmembrane topology of proteins was performed via TMHMM (http://www.cbs.dtu.dk/services/TMHMM/) tool. Antigenicity of the conserved fragments were determined via VaxiJen v2.0 server (http://www.ddg-pharmfac.net/vaxijen/) (Krogh et al., 2001; Doytchinova and Flower, 2007b). The foremost potent antigenic epitopes were chosen and permitted for advance investigation.

### 5. Population coverage, allergenicity and toxicity analysis of predicted epitopes

HLA dispersion changes in diverse geographic spaces and ethnic social orders in the whole earth. IEDB population coverage calculation server (http://tools.iedb.org/population/) was used for population coverage study (Vita et al., 2014). In addition, four servers named AllergenFP (http://ddg-pharmfac.net/AllergenFP/), AllerTOP (http://www.ddg-pharmfac.net/AllerTop/), Allermatch (http://www.allermatch.org/ allermatch.py/form) and Allergen Online (http://www.allergenonline.org) were utilized to foresee the allergenicity of the proposed epitopes for immunization development (Dimitrov et al., 2014; Dimitrov et al., 2013; Fiers et al., 2004) whereas toxicity was predicted by using ToxinPred tool (http://crdd.osdd.net/raghava/toxinpred/) (Gupta et al., 2013).

### 6. Conservancy analysis of MHC restricted alleles

Conservancy investigation server (http://tools.iedb.org/conservancy/) was chosen for conservancy investigation at IEDB server. In this case, protein BLAST was done with antigenic proteins which was picked from the NCBI database.

### 7. Identification of B-Cell epitope

B-Cell epitopes were identified by using IEDB server. In this case, 3 several algorithms are used and those are: Bepipred Linear Epitope Prediction 2.0, Emini surface accessibility prediction and Kolaskar and Tongaonkar antigenicity scale (Jespersen et al., 2017; Emini et al., 1985; Kolaskar and Tongaonkar, 1990).

### 8. Construction of vaccine molecules

Three vaccine constructs (i.e. V1, V2 and V3) were built. Different adjuvents like beta defensin (a 45 mer peptide); HABA protein (accession number: AGV15514.1; *M. tuberculosis*) and L7/L12 ribosomal protein were used in this purpose after the best CTL epitopes, best HTL epitopes and BCL epitopes individually(Rana and Akhter, 2016). Other sequence like PADRE as well as diverse linker like GGGS, EAAAK, KK and GPGPG were utilized too for developing of potential vaccine against monkeypox virus.

### 9. Allergenicity, antigenicity and solubility prediction of various vaccine constructs

Four tools named AllergenFP, AllerTOP, Allermatch and Allergen Online were utilizedfor determining the non-allergic nature of the constructed vaccine. To determine the most potent vaccine, antigenicity and the solubility of the suggested vaccine were determined via VaxiJen v2.0 and via Proso II tool respectively (Doytchinova and Flower,2007b; Smialowski et al., 2006).

### 10. Physicochemical characterization and secondary structure analysis

ProtParam, a tool provided by ExPASy server (Hasan et al., 2019b; Azim et al., 2019a) was used to functionally characterize (http://expasy.org/cgibin/protpraram) (Gasteiger et al., 2003) the vaccine proteins. Isoelectric pH, aliphatic index, molecular weight, instability index, estimat half-life, hydropathicity, GRAVY values, and other physicochemical characteristics were studied. Through GOR4 secondary structure prediction method, alpha helix, beta sheet and coil structure of the vaccine constructs were determined by the Prabi tool (https://npsa-prabi.ibcp.fr/).

### 11. Structure predictio, validation and disulfide engineering of vaccine constructs

RaptorX server was used for constructing 3D model of vaccine. This step is necessary for knowing the percentage of sameness between template structure and target protein and those were collected from RCSB PDB (Peng and Xu, 2011). Vaccine structure was validated by Ramachandran plot assessment at RAMPAGE (Hasan et al., 2015). Disulfide bonds was designed by DbD2 server for anticipated vaccine constructs (Craig and Dombkowski, 2013).

### 12. Protein-protein docking

Patchdock tool was used for molecular docking between HLA alleles and the predicted vaccine constructs. The tool leveled the docked compounds based on complementarity score, ACE (Atomic Contact Energy) and approximate interface area of the compound. On the basis of free binding Energy, results showed that V1 vaccine was the superior. V1 construct also docked with various human immune receptors (TLR 2, TLR 4,TLR 9). Highest binding affinity between vaccine construct and receptor were detected by the lowest binding energy of the complexes.

### 13. Codon adaptation and in silico cloning

Codon adaptation was carried out via JCAT tool to speed up the expression of construct V1 in *E. coli* strain K12. In this case, various restriction enzymes (i.e. BglI and BglII), prokaryote ribosome-binding site and Rho independent transcription termination were kept away from the work (Grote et al., 2005).Than mRNA sequence of constructed V1 vaccine was conjugated with BglI and BglII restriction site at the C-terminal and N-terminal sites respectively. In this case, SnapGene was used for the cloning purpose (Solanki & Tiwari, 2018).

## Result

### 1. Retrieval of protein sequences and antigenicity screening

The full viral proteome of monkeypox virus (Proteome ID: UP000101269) was collected from UniProtKB (https://www.uniprot.org/uniprot/?query+database). Most potent antigenic protein was found from the VaxiJen server. Among three viral proteins, cell surface binding protein (Accession ID: Q8V4Y0.) and Poxin-Schlafen (Accession ID: Q8V4S4) and Envelope protein A28 homolog (Accession ID: Q8V4U9) were identified, having better immunogenic potential with total prediction score of 0.5311, 0.4753 and 0.6212 respectively. Various types of physiochemical properties of proteins are given in Table 01.

**Table 01:** Protparam analysis of selectet protein

| **protein** | **Accession ID** | **Molecular weight** | **Instability index** | **Half-life** | **Theoretical pI** | **No. of Amino acids** | **Total No. of Atoms** | **Extinction co-efficient** |
| --- | --- | --- | --- | --- | --- | --- | --- | --- |
| Cell surface binding protein | Q8V4Y0 | 35278.02 | 44.95 | >10 hours | 7.77 | 304 | 4953 | 56270 |
| Envelope protein A28 homolog | Q8V4U9 | 16403.68 | 29.97 | >10 hours | 6.54 | 146 | 2286 | 14440 |
| Poxin-Schlafen | Q8V4S4 | 57453.78 | 39.03 | >10 hours | 7.58 | 503 | 8029 | 76210 |

### 2. Retrieval of homologous protein sets and identification of conserved regions

Three several homologous protein of monkeypox virus were created after BLASTp via NCBI BLAST server. Different homologous protein sets for each protein were generated after BLASTp search using NCBI BLAST tools. A total of 6, 4 and 10 conserved pieces were found among the cell surface binding protein, envelope protein A28 homolog and poxin-Schlafen protein respectively (Table 02).

**Table 02:** Identified conserved regions among different homologous protein sets of Cell surface binding protein, Envelope protein A28 homolog and Poxin-Schlafen

| Protein | Conserved region | Vexigen | Topology |
| --- | --- | --- | --- |
| Cell surface binding protein | MPQQLSPINIETKKAISD | 0.9972 | outside |
|  | RLKTLDIHYNESKPTTIQNTGKLVRINFKGGYISGGFLPNEYV<br>LSTIHIYWGKEDDYGSNHLIDVYKYSGEINLVHWNKKKYSS<br>YEE | 0.7807 | outside |
|  | KKHDDGIIIIAIFLQVSDHKNVYFQKIVNQLDSIRSANMSAPF<br>DSVFYLDNLLPSTLDYFTYLGTINHSADA | 0.4468 | outside |
|  | WIIFPTPINIHSQDLSKFRLLSSSNHEGKP | -0.0067 | outside |
|  | YITENYRNPYKLNDDTQVYYSGEIIRAATTSPVRENYFMKW<br>LSDLR | 0.1645 | inside |
|  | CFSYYQKYIEGNKTFIIAIVFVILT | 0.6078 | outside |
| Envelope protein A28 homolog | MNSLSIFFIVVATAAVCLLFIQSYSIYENYGNIKEFNATHAAF<br>EYSKSIGGTPALDRRVQDVND | 0.6333 | inside |
|  | ISDVKQKWRCVVYPGNF | 0.9445 | inside |
|  | SASIFGFQAEVGPNNNT | 0.7511 | outside |
|  | SIRKFNTMRQCIDFTFSDVINIDIYNPCIANINTECQFLKSV<br>L | 0.4689 | inside |
| Poxin-Schlafen | MFYAHAFFGGYDENLHAFP | 0.2036 | outside |
|  | ISSTVANDVRKYSVVSVINKKYNIVKNKYMWCNSQVNKR<br>YIGALLPMFECNEYLQIGDPIHDLEGNQISIVTYRHKNYYAL<br>SGIGYESLDLCL | 0.6140 | outside |
|  | GVGIH HHVLETGNAVYGKVQHEYSTIKEKAKEMNA | 0.7427 | outside |
|  | KPGPIIDYHVVWIGDCVCQVTTVDVHGKEIMRMRFKRGAVL<br>PIPNLVKVKVGEENDTINLSTSISALLNSGGGTIEVTSKEERV<br>DYVLMKRLESIHHLWSVVYDHLNVVNGEERCY | 0.5687 | outside |
|  | TVKTNLYMKTMGACL | 0.4971 | inside |
|  | MDSMEALEYSELKESGGRSPPELQKFEYPDGVKDTESIE<br>RLAEFFNRSELQAGESVKFG | 0.2660 | outside |
|  | SINVKHTSVSAKQLRTRIR | 1.3666 | inside |
|  | QLPSILSSFANTKGGYLFIGVDNNTHKVIGFTVGHDLKLV<br>E | 0.5208 | outside |
|  | DIEKYIQKLPVVHFCKKKEDIKYACRFI | 0.2575 | inside |
|  | IKVERCCCAVFADWPESWYMDTSGSMKKYSPDEWVSHIKF | 0.2159 | inside |

### 3. Antigenicity prediction and transmembrane topology analysis of the conserved fragments

4,4,5 conserved framents were selected from cell surface binding protein, envelope protein A28 homolog and poxin-Schlafen proteins respectively because of their VaxiJen level is upper than their default threshold level (Table 02). On the other hand, transmembrane topology server showed that, from the immunogenic conserved sequences 5, 1 and 6 sequences from the corresponding proteins fulfilled the property of exomembrane characteristics (Table 02).

### 4. Prediction of T-cell epitope, transmembrane topology and antigenicity screening

A large number of epitops were created from the conserved sequences which could bind with highest number of HLA cells with maximum binding affinity. After analysis their antigenicity score and transmembrane topology, top epitopes of the selected three proteins were screened (Table 03). Epitopes with a positive higher value of immunogenicity showed potential to evoket effective T-cell response.

**Table 03:**
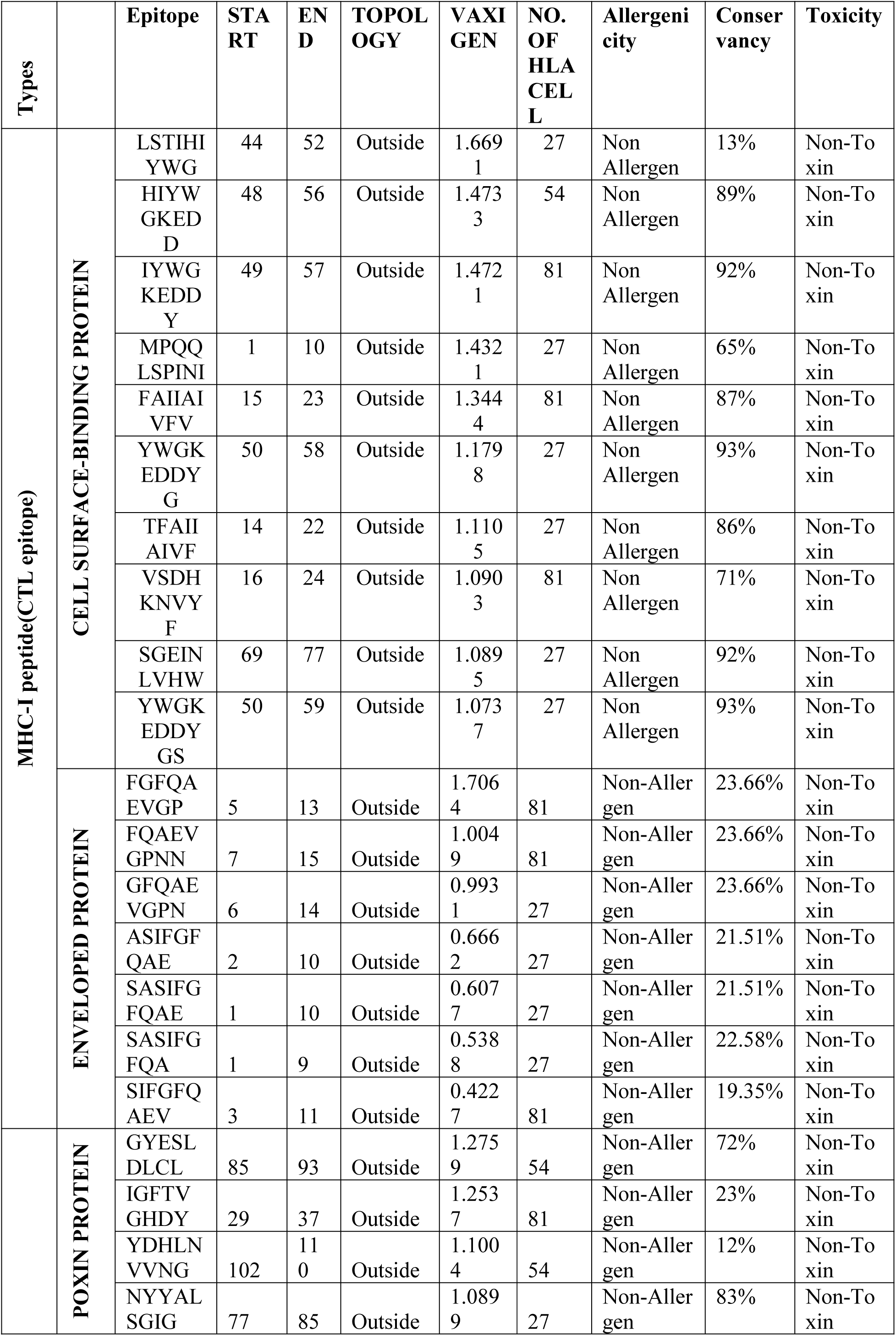
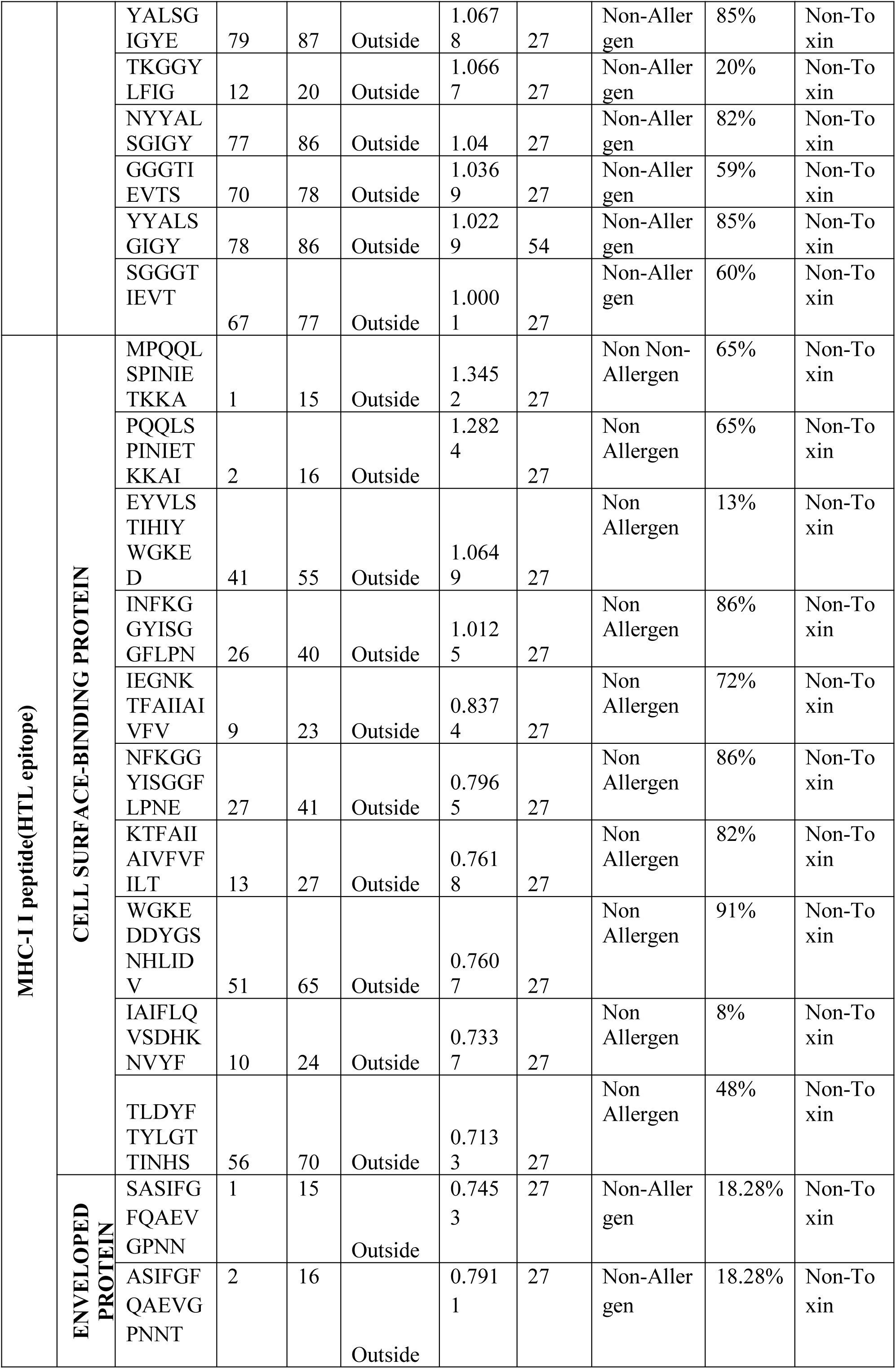
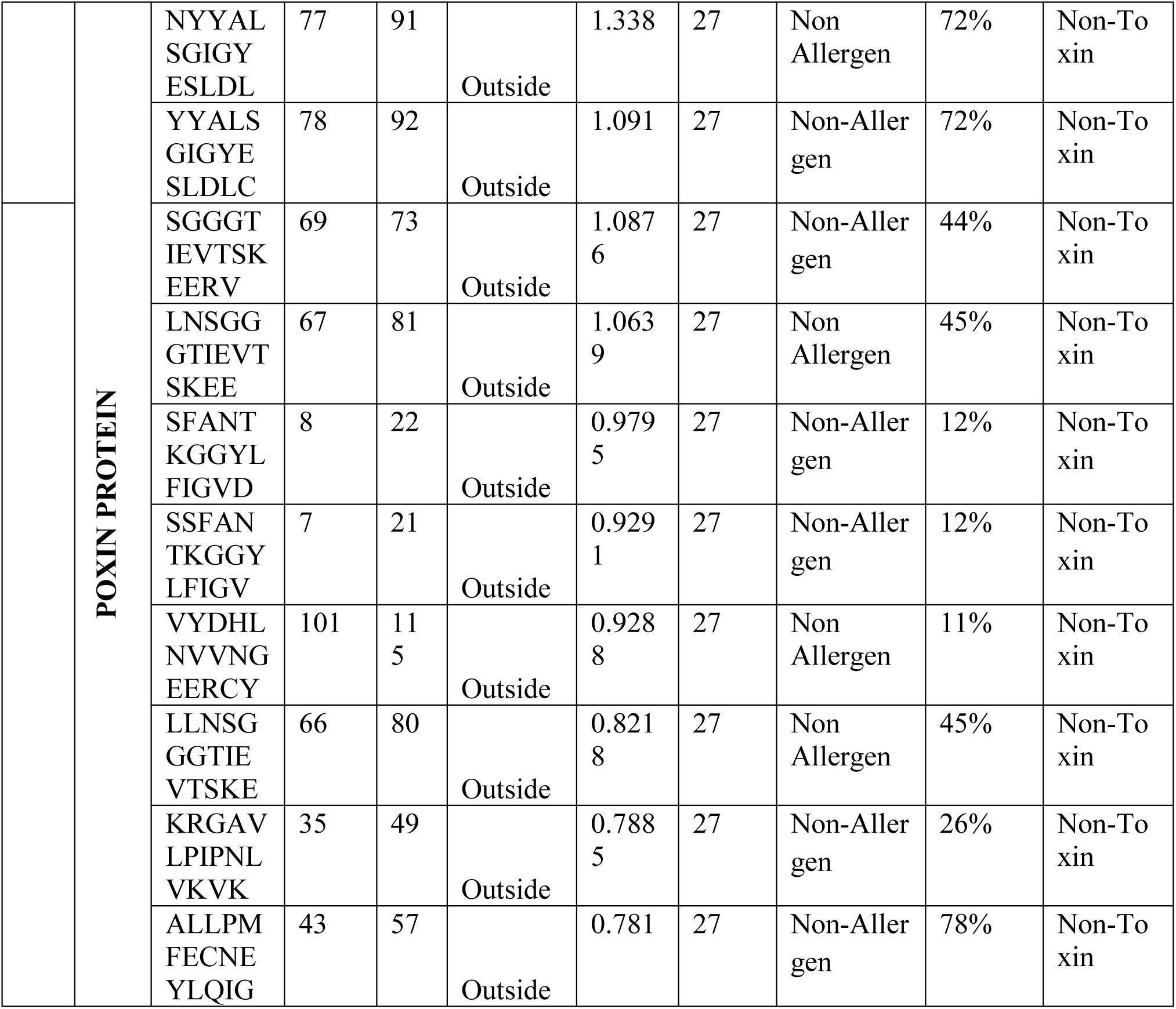
Predicted T-cell (CTL and HTL) epitope of Cell surface binding protein, Envelope protein A28 homolog and Poxin-Schlafen

### 5. Population coverage, allergenicity and toxicity analysis of predicted epitopes

Popultion coverage were done for all three proteins with their both MHC class I and MHC class II. From the screening, populationof thevarious geographic areas can be covered by the predicted T-cell epitopes.results of population coverage of three several viral proteins are shown in Figure 2. AllerTOP, AllergenFP, Allergen online, Allermatch servers were used for allergenicity screening. Non-allergen epitopes for human were selected (Table 03). Allergen and toxic epitopes were removed from the predicted list (Table 03).

**Figure 02:**
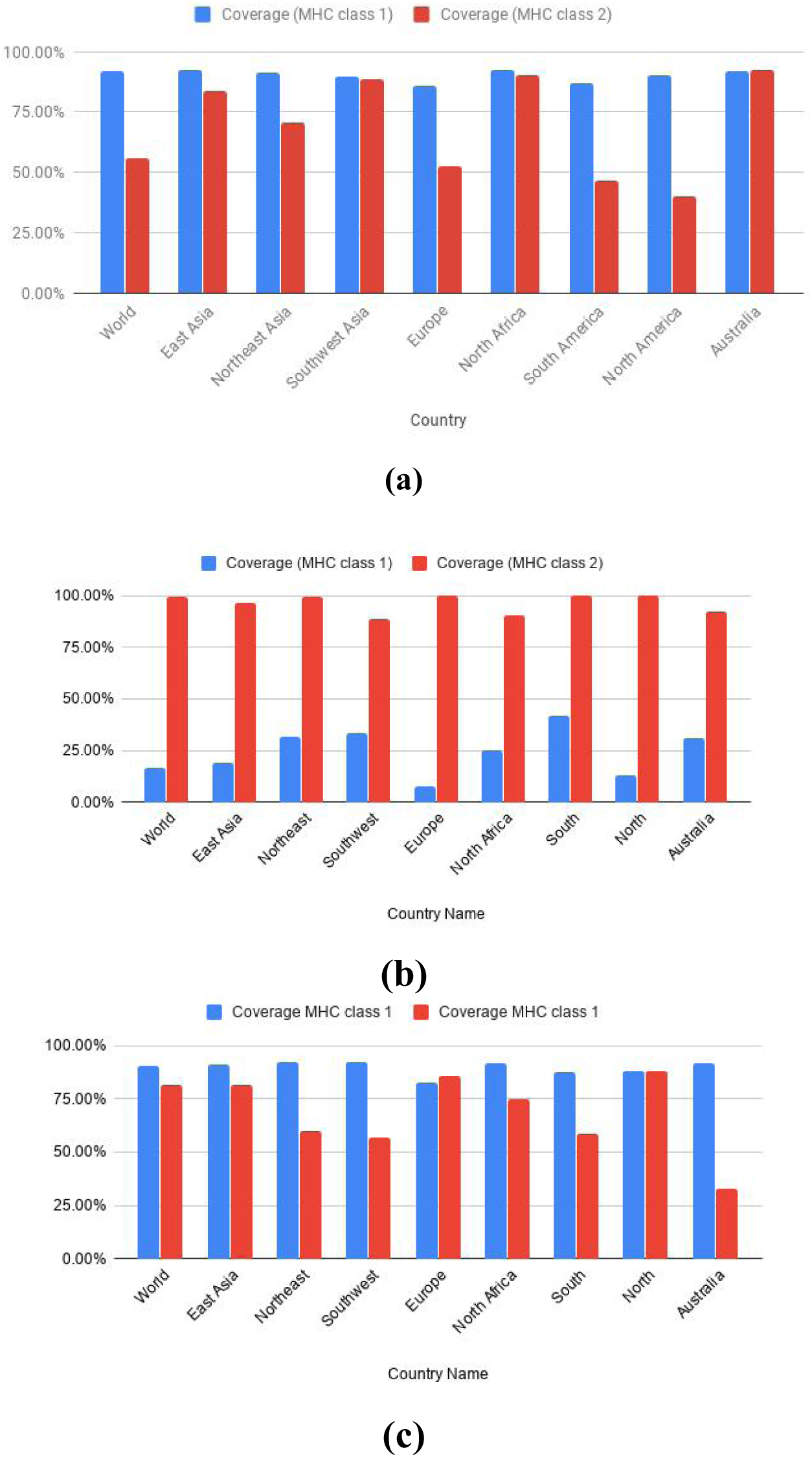
Population coverage analysis of predicted T-cell epitopes (MHC-I and MHC-II peptides) for Cell surface binding protein (a); Envelope protein (b) and Poxin-Schlafen (c).

### 6. Epitope conservancy analysis

Cell surface binding protein, poxin-Schlafen and envelope protein were found to be exceedingly moderated inside diverse strains of monkeypox infection with most noteworthy conservancy level of 93% (Table 01). The rate of conservancy chowed that, epitopes were organically noteworthy.

### 7. Identification of B-Cell epitope

Three algorithms (i.e. Bepipred Linear Epitope prediction, Emini Surface Accessibility, Kolaskar & Tongaonkar Antigenicity prediction) were used for screeninf of B-cell epitopes of cell surface binding protein, poxin-Schlafen and envelope protein from IEDB. Further screening of selected epitopes were done by analysis their antigenicity scoring and allergenicity (Table 04).

**Table 04:** Allergenicity assessment antigenicity investigation of the anticipated B-cell epitopes

| **Protein** | **Algorithm** | **Top Epitope** | **Allergenicity** | **VaxiJen score** |
| --- | --- | --- | --- | --- |
| Surface-Binding Protein | Linear epitope | KYSGEINLVHWNKKKYS<br>SYEEKKHDDG | Non-allergen | 0.8238 |
|  | Surface accessibility | HYNESKPTTI | Non-allergen | 0.5181 |
|  | Antigenicity | HKNVYFQKIVNQL | Non-allergen | -0.0446 |
| Enveloped protein | Linear epitope | GFQAEVG | Non-allergen | 0.5058 |
|  | Surface accessibility | VGPNTT | Non-allergen | 1.2433 |
|  | Antigenicity | IFGFQAE | Non-allergen | 0.7731 |
| Poxin Protein | Linear epitope | DYHVVWIGDCVCQVTTVD | Non-allergen | 0.3489 |
|  | Surface accessibility | KVQHEYSTIKEKAKEMN<br>AK | Non-allergen | 0.8280 |
|  | Antigenicity | GAVLPIPNLVKVK | Non-allergen | 0.4370 |

### 8. Vaccine molecule construction

Total three vaccines (V1, V2 and V3) were constructed (Table 05). Each vaccine construct contain a protein adjuvent, T-cell, B-cell epitopes and epitope’s linkers. PADRE sequence was added to increase the potency and efficacy of the constructed vaccine.The designed vaccine constructs V1, V2 and V3 were 323, 407 and 348 residues long.

**Table 05:**
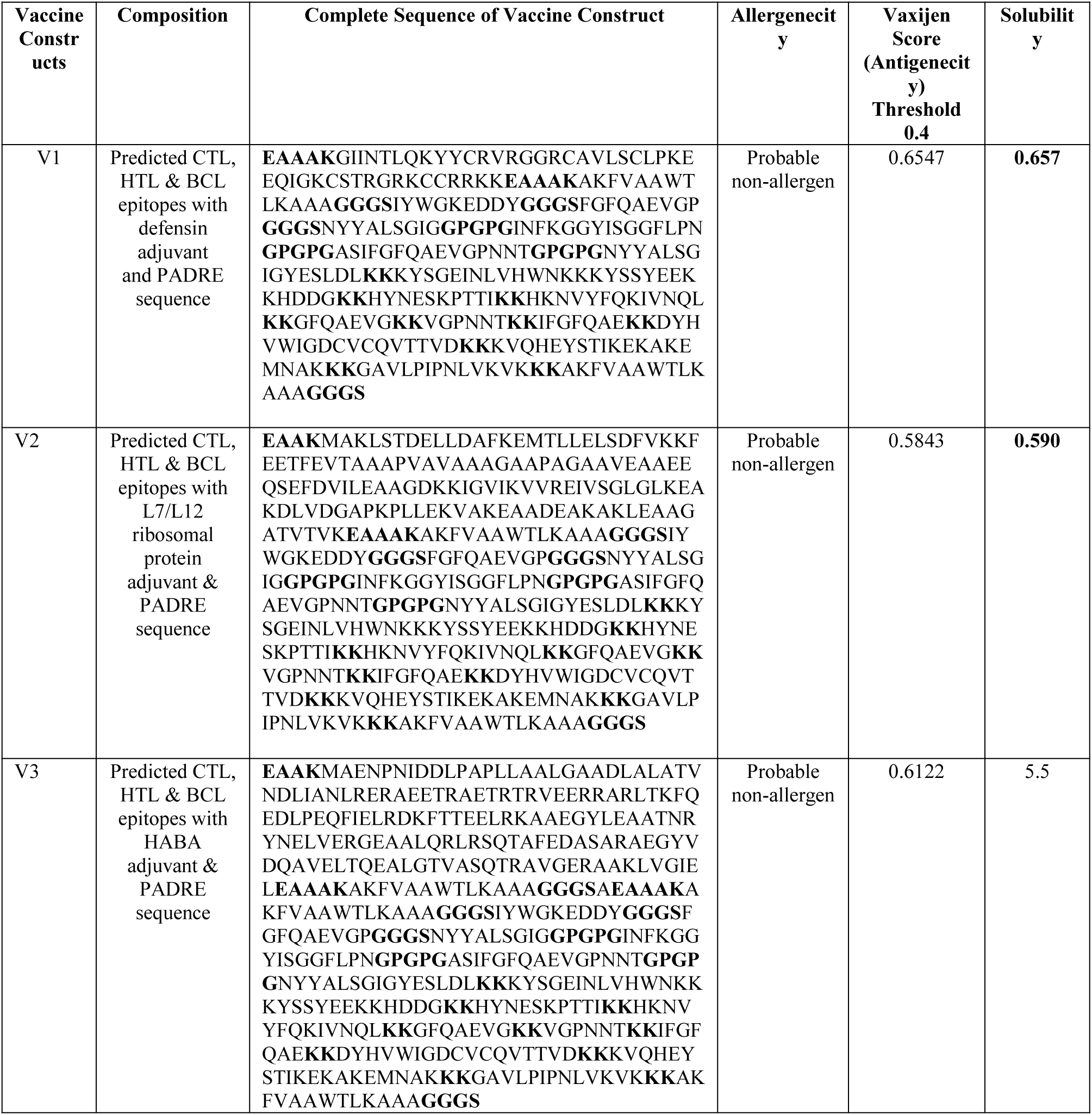
Allergenicity, antienicity and solubility prediction of the designed vaccine constructs

### 9. Allergenicity, antigenicity and solubility prediction of different vaccine constructs

All 3 constructs (V1, V2 and V3) were non-allergenic (Table 05). All of them, V1 showed the best result with solubility score (0.657) (Figure 3) and antigenicity (0.6547).

**Figure 03:**
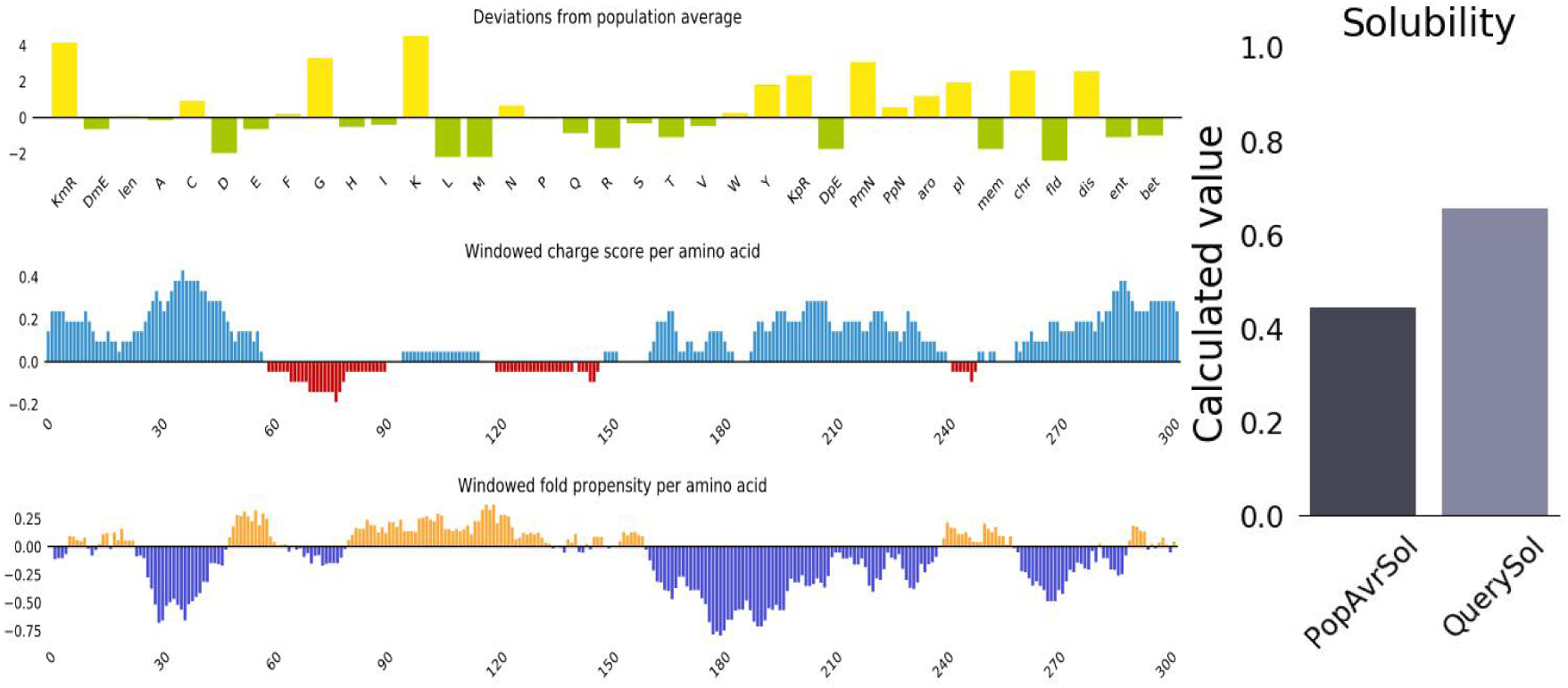
Solubility prediction of designed superior vaccine V1 using via Protein-sol server.

### 10. Physicochemical characterization and secondary structure analysis

Physicochemical properties of V1 construct were analyzed through ProtParam tool. Molecular weight of V1 construct showed superior immunity and it was scored 34.879 kDa. The extinction coefficient was 52830 at 0.1% absorption, assuming all cysteine residues are lowed. The theoretical pI 9.75 recommended that the protein will have net negative charge which is higher than this pI. The approximated half-life was decided to be one hour within mammalian reticulocytes in vitro while greater than ten hours in *E. coli* in vivo. Thermostability and hydrophilic nature of the protein were presented by the aliphatic index and GRAVY value which were 62.26 and −0.619 respectively. Secondary structure of the predictedt V1 vaccine construct confirmed to have 27.24% alpha helix, 23.84% sheet and 48.92% coil structure.

### 11. Prediction, validation and disulfide engineering of vaccine constructs

V1 vaccine constructs was composed by using RaptorX server (Figure 04). Ramachandran plot analysis was used during validation process where 90.9% residues were in the favored, 7.2% residues in the allowed and 1.9% residues in the outlier region (Figure 04). 3D structure of V2, V3 vaccine were constructed (Figure 05). There were 33 pairs of amino acid residue were recognized which have the potentiality to create disulfide bond by DbD2 server. After analysis chi3 and B-factor parameter of residue pairs on the basis of energy, only two pairs (GLY 21-ARG47 and LYS245-SER250) fulfill the property for disulfide bond development which were changed with cysteine. The value of energy was considered less than 2.5 and chi3 value for the residue screening was between –87 to +97 (Figure 06).

**Figure 04:**
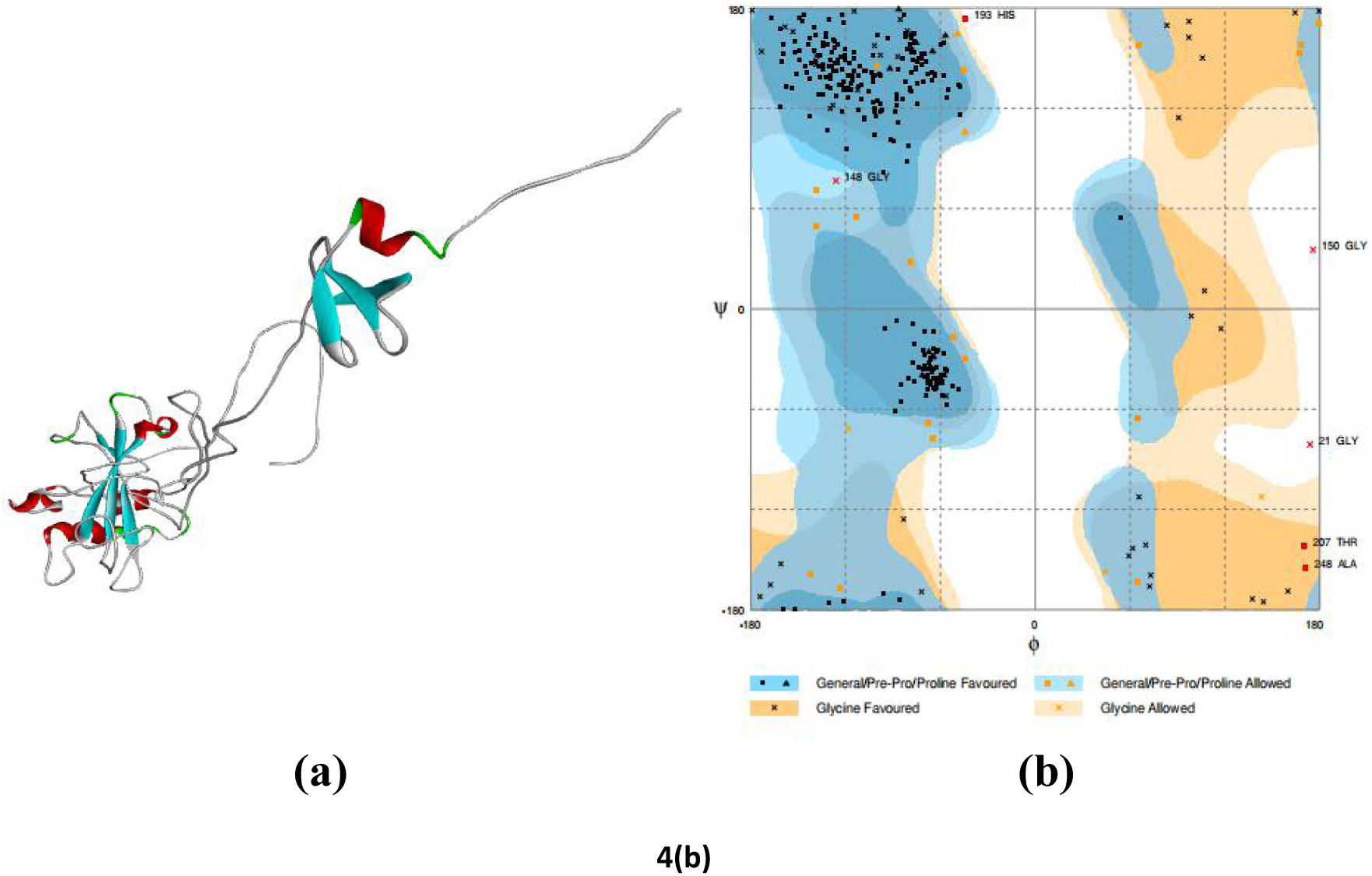
Tertiary structure of vaccine construct V1(a) and validation of it’s tertiary structure via by Ramachandran plot analysis (b).

**Figure 05:**
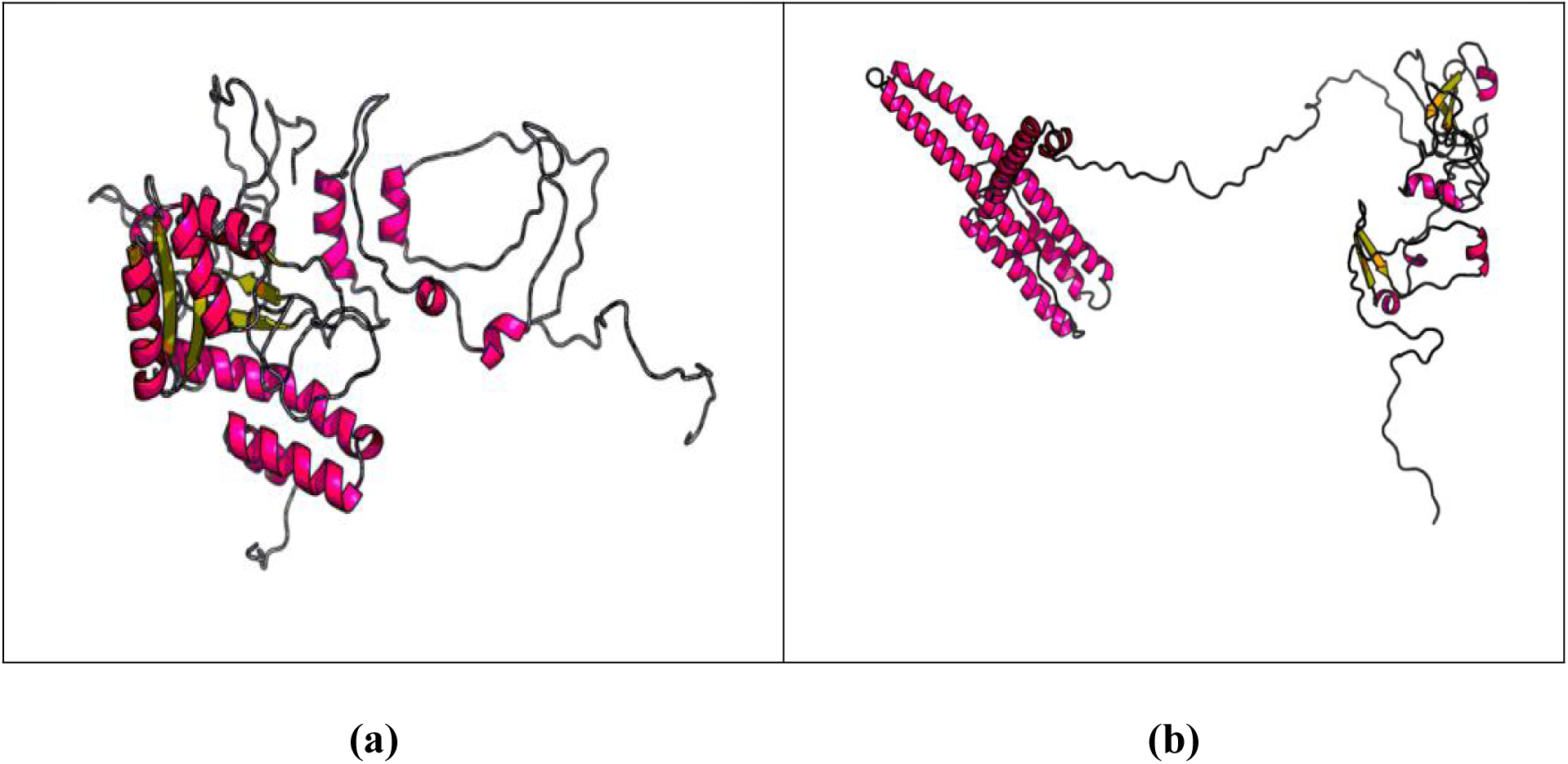
Tertiary structure of V2 (a) and V3 (b)vaccine constructs.

**Figure 06:**
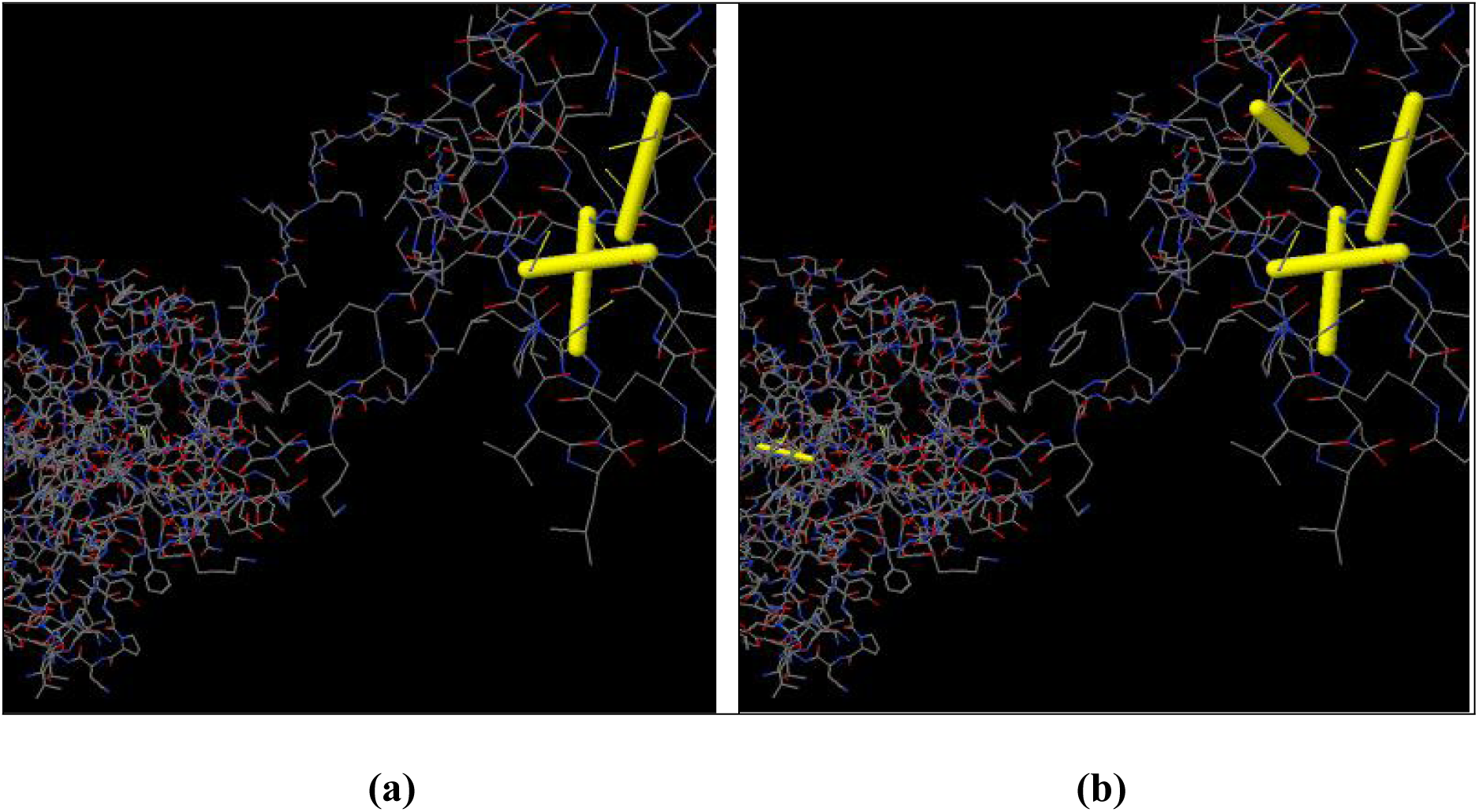
Disulfied engineering of predicted superior vaccine construct V1; original (a) and mutant (b).

### 12 Protein-protein docking

The affinity between HLA alleles and the constructed vaccines (V1, V2 and V3) were analyzed using molecular docking via Patchdock tool. The affinity between V1 construct and human immune receptors were determined also (Figure 07). The server leveled the docked compounds based on complementarity score, Atomic Contact Energy (ACE) and approximate interface area of the complex. All of them, V1 vaccine construct showed the superior in terms of free binding Energy (Table 07). Highest binding affinity between vaccine construct and receptor were indicated by ther lowest binding energy of the complexes.

**Figure 7:**
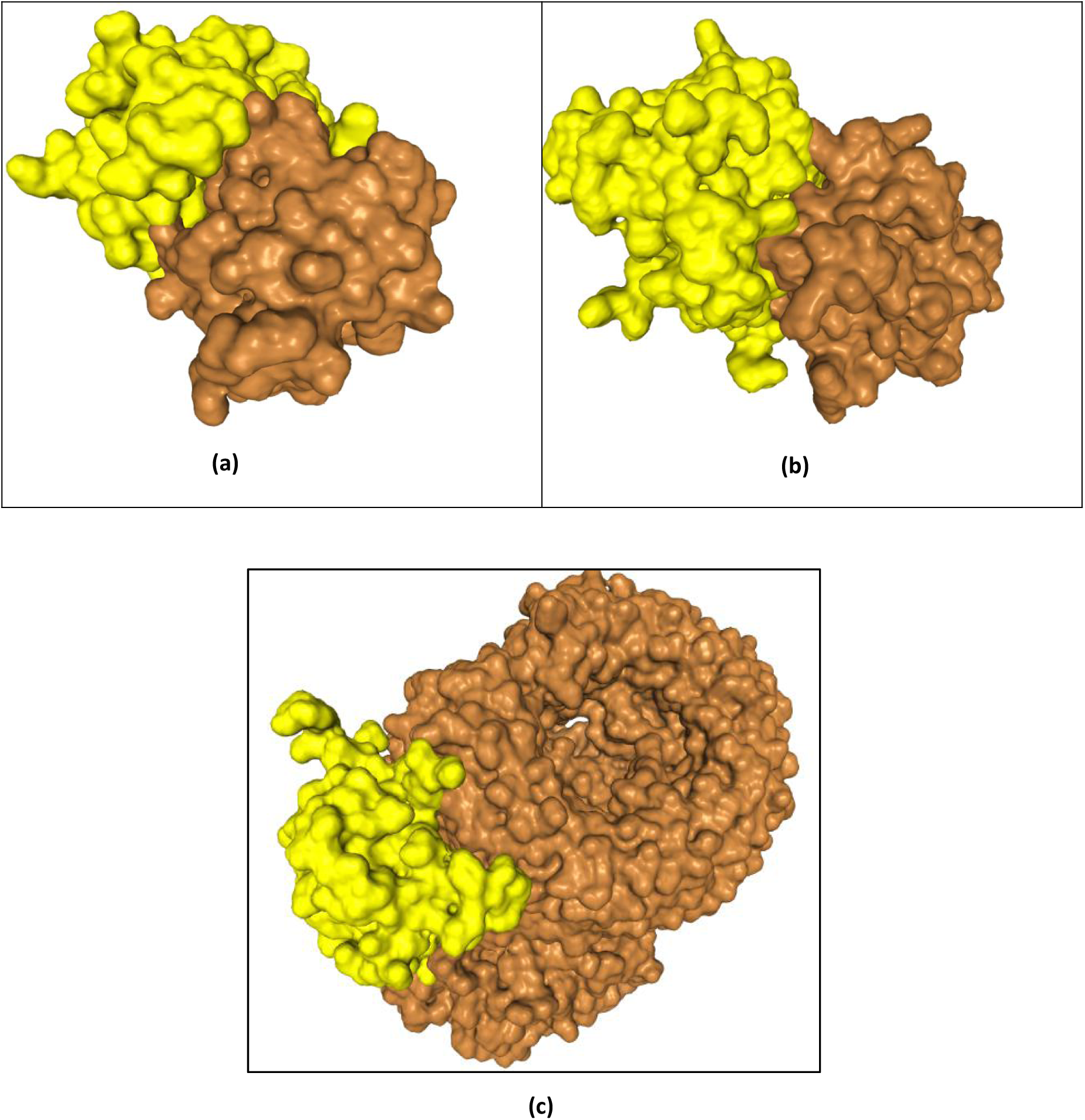
Molecular docking between vaccine V1 with MyD88 (a), TLR2 (b) and TLR8 (c).

### 13 Codon adaptation and in silico cloning

The Codon Adaptation Index for the optimized codons of construct V1 was 0.967 determined via JCAT server. The GC content of the adapted codons was also significant (46.23%). An insert of 986 bp was obtained which lacked restriction sites for BglII and BglI ensuring, thus ensuring safety for cloning purpose. The codons were inserted into pET28a(+) vector along with BglII and BglI restriction sites and a clone of 4564 base pair was produced (Figure 08).

**Figure 08:**
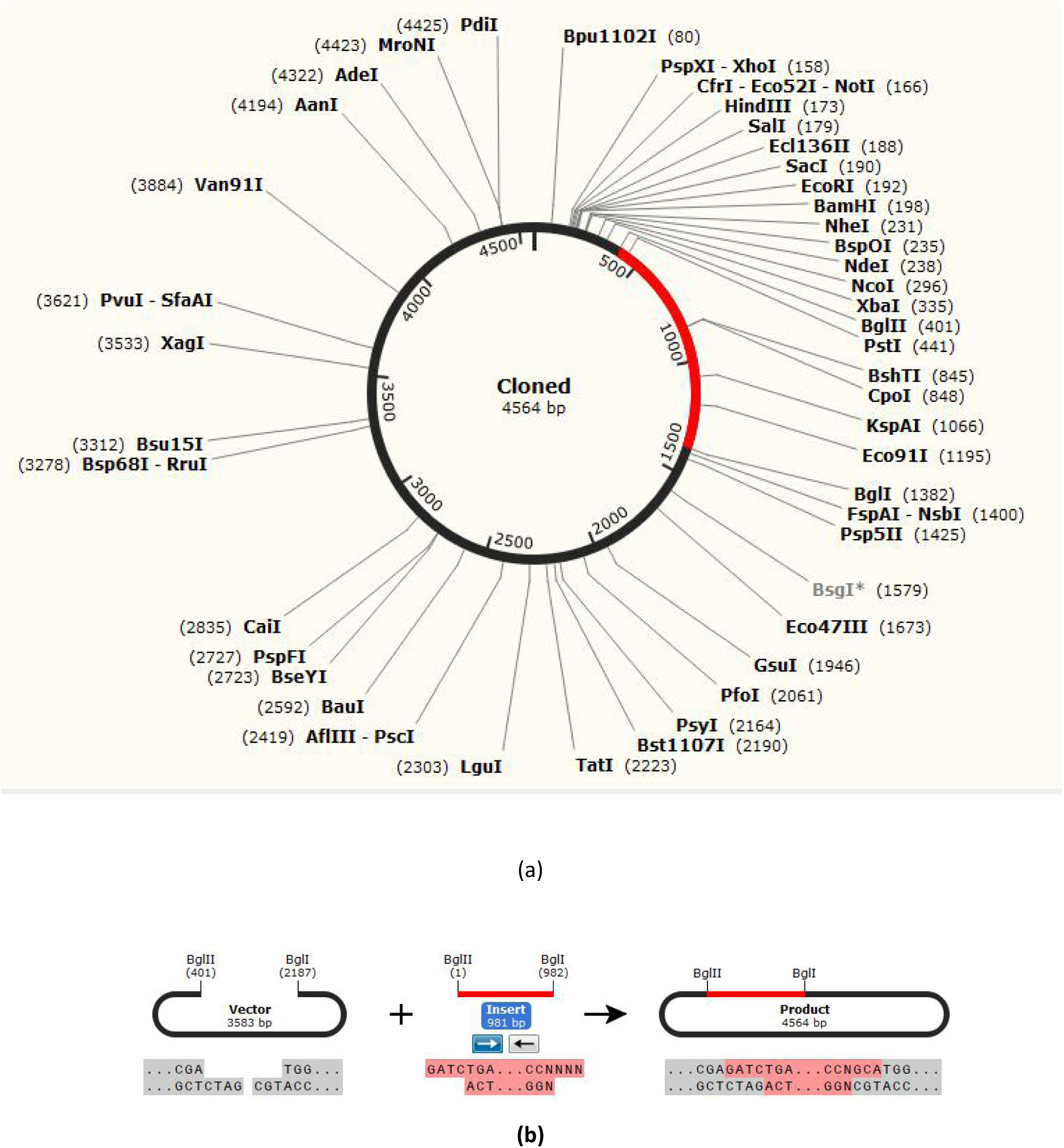
Restriction digestion and *in silico* cloning of the gene sequence of final construct V1 into pET28a(+) expression vector. Target sequence was inserted between BglII (401) and BGlI (2187).

## Discussion

Monkeypox virus (MPX) has recently focused global media, political and scientific attention after the identification in the United Kingdom (UK) in September 2018 of 3 separate patients diagnosed with monkeypox (PHE, 2018). Though it causes disease in humans with symptoms similar, but less severe, to those previously seen in smallpox (WHO, 2019), the extremely infectious virus is often responsible for severe systemic lesional disease in humans and noteworthy auxiliary spread of infection inside near communities. Along these lines, it is imperative to require preventive measures against it. Generally, live-attenuated or inactivated forms of microbial pathogens have been utilized for induction of antigen-specific responses that prevent the host against secondary infections (Thompson & Staats, 2011). But only 10 of a few hundred proteins are used for vaccine development and it depends on the microbes (Li et al., 2014). However, all of the protein is not necessary for protective immunity, whereas only few selective proteins are used (Tesh et al., 2002). On the other hand, those unnecessary proteins are responsible for allergenic reactions. So it is crying need to remove all of those protein from the vaccine formulations (Petrovsky & Aguilar, 2004). There were no licensed therapies to treat human monkeypox viral infection (Pal et al., 2017) until 2019. Few days ago, FDA accepted a vaccine called JYNNEOS which is a non-replicating and live vaccine for the avoidance of monkeypox virus. Be that as it may, no security concerns that would require a Medication Guide were determined (ANALGESICS, 2019). The foremost common (>10%) antagonistic responses related with JYNNEOS were infusion location reactions. Itching, swelling, pain, induration, redness, are the common side effects of this vaccine. There are also some systemic adverse reactions also which are headache, nausea, muscle pain, fatigue, chills and myalgia. Genuine antagonistic responses were detailed in 0.05% of subjects who gotten JYNNEOS and included sarcoidosis, throat tightness, Crohn’s malady and extraocular muscle paresis. Cardiac unfavorable responses of extraordinary intrigued were detailed in 0.08% of subjects who gotten JYNNEOS and added electrocardiogram T wave inversion, electrocardiogram T wave abnormal, electrocardiogram ST segment elevation, palpitation and electrocardiogram abnorma (Bavarian-nordic, 2019). ST-246 is an anti orthopoxvirus compound that extremely potent, inhibiting orthopoxvirus replication in vitro at nM concentrations (Duraffour et al., 2007; Quenelle et al., 2007; Yang et al., 2005). after injection of ST-246, lesions are created and weight is also lost (Grosenbach et al., 2008). For these reasons, it is necessary to develop subunit vaccines which contain one or few selective proteins of pathogens in vaccine construction for induction of protective immunity (Petrovsky & Aguilar, 2004; Thompson et al., 2011). However, this can create an attention towards “peptide vaccines”. This vaccine contains only epitopes able to induce positive, desirable T cell and B cell mediated immune response (Sesardic, 1993). To activate humoral and cellular responses, these peptides are considered adecuate for activation (Bijker et al., 2007; Lin et al., 2013), whereas killing allergenic and/or reactogenic reactions.

Ordinary approaches for vaccine advancement are not continuously attainable as pathogens are some of the time troublesome to develop and in a few occurrences to constrict coming about in unpleasant or antagonistic immune reactions (Purcell et al., 2007; Azim et al., 2019b). Propels in genomics have altered the concepts and presently, it is conceivable to utilize genomic based approaches to help determination and structure based plan of vaccine candidates (Ahluwalia et al., 2017). Reverse vaccinology, a novel process to combine immunogenetics and immunogenomics with in-silico study has been utilized massively to present modern vaccines (Hasan et al., 2019c; Poland et al., 2009). In this study, we pointed to plan an epitope based peptide vaccine against monkeypox through immunoinformatics procedures by using different bioinformatics database and servers. To the finest of our information such genome based immunoinformatic process has not be used however to create a multiepitope vaccine with wide run viability. The full monkeypox virus proteome was collected from UniProtKB and the physiochemical characteristics of the proteins were examined. The VaxiJen tool was utilized to survey the antigenicity of all the recovered protein arrangements in arrange to discover out the foremost potent antigenic protein. Among the three monkeypox proteins, ell surface binding protein, Poxin-Schlafen, Envelope protein A28 homolog were recognized as the finest immunogenic protein candidates and permitted for advance investigation. All the proteins are selected on the basis of antigenicity. Since antigenicity has been utilized to depict both the capacity of the antigen to combine with the antibody conjointly its capacity to induce antibody creation (Rao, 1972). These proteins have necessary function in membrane fusion, recognition by host immune machinery and receptor binding (Oster, 2016). Envelope proteins of monkeypox virus has specificity for adding to human cells (Maa et al., 1990). Poxin is a nuclease enzyme that is conserved in mammalian poxviruses (Eaglesham et al., 2019). Poxin is preserved in mostvirus within the class Orthopoxvirus, counting the human pathogens monkeypox infection and cowpox infection, and is in some cases combined to an extra C-terminal space to have homology with human schlafen proteins (Liu et al., 2018). Two eminent exemptions to preservation of poxin action are Variola major infection, the causative specialist of smallpox malady, and the adjusted vaccinia Ankara (MVA) immunization strain, which both appear inactivation of poxin (Eaglesham et al., 2019). Preservation of poxins between mammalian infections and creepy crawlies reaffirms the hereditary roots of 2′,3′-cGAMP-dependent flagging in creatures, and underscores the wide extend of host–virus clashes that drive modern components of intrinsic safe reconnaissance and avoidance (Kranzusch et al., 2015). To guarantee defensive reaction against a longer run of infection strains for a longer period, the selected epitopes must stay within the profoundly conserved locale. For this, only the conserved sequences are used for epitope recognition. In this study, we believe that, we can create a universally effective and more acceptable vaccine constructs. Assurance of potent T cell epitopes could be a urgent step amid vaccine design, since T-cells play the key part in antibody generation through antigen introduction. T cell epitopes identification is the major step at the time of vaccine design. Moreover, T-cells play the key role in antibody production through antigen presentation (Amorim et al., 2016). IEDB T cell epitope prediction server was used for predicting both MHC class I (CTL) and MHC class II restricted (HTL) epitopes. Transmembrane topology screening, allergenicity pattern, antigenicity scoring, toxicity profile and conservancy analysis were used for selecting top potent epitopes.

The proposed protein appeared a high aggregate population coverage in most of the geographic rlocales counting 92.51% (MHC-I) and 92.49% (MHC-II) coverage in North Africa and Australia individually (Fig. 2). The top epitopes were highly conserved among different viral strains extending from 8% to 91% (Table 3). furthermore, the selected CTL and HTL epitopes were checked for their capacity to tie with MHC-I and MHC-II alleles. B-cell epitopes can boost neutralizing-antibody responses in the host. To realize defensive immunity against monkeypox virus, B-cell (BCL) epitopes were recognized utilizing three distinctive algorithms (i.e. Linear Epitope, Surface Accessibility, Antigenicity) from IEDB (Table 4). Carrier molecules were needed to add adjuvanting and chemical stability to the epitopes, because the size of epitoe is tiny and have weak immunogenicity (Aguilar & Rodriguez, 2007; Purcel et al., 2007). The selected CTL, HTL and BCL epitopes were added using various linker and adjuvants to design the final constructs. Than the efficiency and safety were recognized (Table 5). It was found that, V1 is the most potent vaccine construct after analysis allergenicity, physicochemical characteristics, hydrophobicity and immunogenicity score. Tertiary structure of the vaccine constructs were determined to strengthen our selection. Moreover, interactions between various HLA molecules (i.e. HLA-DRB1*03:01, HLA-DRB5*01:01, HLA-DRB1*01:01, HLA-DRB3*01:01, HLA-DRB1*04:01 and HLA-DRB3*02:02) and designed vaccine constructs were identified (Table 6). On the basis of free binding energy, V1 vaccine construct was found to be the best. Besides, docking examination was too performed to investigate the binding affinity of develop V1 and distinctive human immune receptors (TLR 2, MyD88, TLR 8) to assess the adequacy of utilized adjuvants (Table 6). Those human immune receptors were identified for the cellular receptor of the monkeypox virus (Schleimann et al., 2019; Xu et al., 2000; Hutchens et al., 2008). The toll-like receptor 8 (TLR8), is found essentially in endosomes of plasmacytoid dendritic cells and B cells where it identifies non-methylated cytosine-guanine DNA motifs (Kawai & Akira, 2011; Kapp et al., 2014). At last, the develop V1 was switch deciphered and embedded inside pET28a(+) vector for heterologous expression in *E. coli* srain K12. Due to the empowering discoveries of the study, we recommend assist wet lab based investigation utilizing show creatures for the exploratory approval of our anticipated vaccine candidates. Additionally, novel antigen conveyance frameworks such as Nano-delivery stages might upgrade the viability of the proposed antibody (Hojo 2014; Trovato and Berardinis, 2015).

**Table 06:** Docking scores of vaccine constracts with different HLA alleles i. e. HLA-DRB1*03:01 (1a6a), HLA-DRB5*01:01 (1h15), HLA-DRB1*01:01 (2fse), HLA-DRB3*01:01 (2q6w), HLA-DRB1*04:01(2seb) and HLA-DRB3*02:02 (3c5j) and TLR 2, TLR 4,TLR 9

| **Vaccine construct** | **HLA alleles PDB ID's** | **Global Energy** | **Hydrogen bond energy** | **ACE** | **Score** | **Area** |
| --- | --- | --- | --- | --- | --- | --- |
| V1 | 1a6a | -34.21 | -2.33 | -1.21 | 13550 | 1718.30 |
|  | 1h15 | -15.11 | -5.47 | 8.40 | 15484 | 2032.20 |
|  | 2fse | -35.94 | -6.33 | -1.47 | 15238 | 2578.00 |
|  | 2q6w | -34.21 | -2.33 | -1.21 | 13550 | 1718.30 |
|  | 2seb | -26.09 | -2.25 | -2.92 | 15120 | 1819.50 |
|  | 3c5j | -31.61 | -3.93 | -1.27 | 15712 | 2028.90 |
| V2 | 1a6a | -11.98 | -9.71 | -0.70 | 19082 | 2680.10 |
|  | 1h15 | 8.72 | 0.00 | -0.44 | 17680 | 2621.30 |
|  | 2fse | 0.00 | 0.00 | 0.00 | 18078 | 2799.70 |
|  | 2q6w | 3.07 | 0.00 | 3.48 | 17258 | 2302.80 |
|  | 2seb | -19.73 | -4.91 | -0.36 | 21592 | 3036.00 |
|  | 3c5j | -13.29 | -3.60 | 12.14 | 20484 | 3360.20 |
| V3 | 1a6a | 0.30 | -4.84 | 1.42 | 17434 | 2319.20 |
|  | 1h15 | -21.55 | -3.68 | 15.24 | 17008 | 2435.50 |
|  | 2fse | 2.14 | -1.00 | 4.96 | 15978 | 3697.00 |
|  | 2q6w | -23.89 | -0.57 | -5.54 | 19066 | 2660.10 |
|  | 2seb | -30.10 | -2.08 | -6.81 | 16648 | 2406.30 |
|  | 3c5j | -32.99 | -2.06 | 3.20 | 16214 | 2755.50 |
| V1 | 2z5v | -17.68 | -2.79 | 3.34 | 12824 | 1596.90 |
|  | 1fyw | -7.36 | -1.87 | 3.68 | 15136 | 2882.00 |
|  | 5D67 | -32.77 | -6.92 | -0.78 | 18654 | 2631.20 |
|  | 3w3k | -12.24 | -3.65 | 12.22 | 19024 | 3013.00 |

## Conclusion

In-silico bioinformatics study can be considered as a promising technique to accelerate vaccine improvement against exceedingly pathogenic living beings. Within the display think about, such approach was utilized to plan a novel peptide vaccine against the foremost dangerous monkeypox. The consider proposes, the proposed immunization seem invigorate both humoral and cellular intervened resistant reactions and serve as a potential immunization against monkeypox virus. Be that as it may, in vitro and in vivo immunological tests are exceedingly suggested to approve the adequacy of outlined vaccine constructs.

